# Towards the automation of Species Distribution Modelling: The AutoMaxEnt routine

**DOI:** 10.64898/2026.09.22.753478

**Authors:** Gonzalo Albaladejo Robles, Jessica Lee Abbate, David Redding

## Abstract

Species distribution models are widely used for the evaluation and analysis of species spatial distributions and responses to environmental changes. Due to this, there’s a wide diversity of tools and algorithms to perform these types of analyses. However, despite being a mature field of study, SDM still requires a large amount of research intervention and decision-making to produce reliable models. This is a natural consequence of each species unique response to environmental factors, but in practice it hinders the potential of SDMs to be used effectively and with confidence across large sets of species. Here we present the AutoMaxEnt repository, a collection of functions and tools for the automation of maximum entropy species distribution models. AutoMaxEnt offers a flexible environment to model species distribution models, automating most of the data pre-processing and preparation and giving the user the possibility of exploring multiple model scenarios automatically or with control over how those scenarios are generated. AutoMaxent offers model fit control, multiple background points generation algorithms, automatic model and variable selection, and study area configuration, among other parameters. AutoMaxEnt allows for the automation of model fitting, evaluation and model selection across different species, study areas and time periods.

## Introduction

We are facing a global biodiversity crisis, with declines in monitored wildlife populations of 73% over the last 50 years (WWF 2024). This biodiversity loos have been mainly driven by anthropogenic pressures such as land-use change, overexploitation, pollution, and climate change (IPBES 2019). These changes not only affect species and ecosystems, but also the ecological services human communities rely on (IPBES 2019). It is for this reason that global conservation strategies and policies are needed to tackle biodiversity loss. The Sustainable Development Goals, set up by the United Nations (UN DESA. 2025), are deeply interconnected with biodiversity conservation and healthy ecosystems. For example, target 15 explicitly addresses the reduction of habitat degradation and biodiversity loss among its objectives. However, biodiversity conservation and long-term planning require large amounts of information regarding the ecological and spatial requirements of species (e.g Diaz-Delgado et al., 2017; IUCN/SSC, 2013). In most cases, we lack this information (e.g Garcia-Rosello et al., 2023), and statistical approaches are needed to fill the gaps (e.g Bowler et al., 2024).

A fundamental aspect to understand how species respond to environmental changes is to understand how they are spatially distributed and which environmental factors are key in shaping species ranges (e.g Coelho et al., 2023; Anta∼o et al., 2022). Motivated by this, a collection of statistical tools has been used to model species niches and predict current and future species distributions. This collection of methods is usually termed Species Distribution Models (SDM hereafter) (e.g Peterson et al., 2011). SDMs have become essential tools informing management strategies and prioritisation efforts under different scenarios of environmental changes (Guisan et al., 2013; Frans et al., 2021). Despite being widely used in conservation and ecology, there are still obstacles to standardising and applying these methods across many species. Some of these challenges arise from the theoretical basis of these tools. SDMs are largely based on species niche theory (Peterson et al., 2011), and therefore species distributions are going to be characterised by their specific relationships with their surroundings (Peterson et al., 2011). This, in essence, requires each species to be modelled using its own set of environmental variables and model structure to recreate its niche adequately.

This limitation is usually circumvented by modelling the potential, rather than the realised niche of species (e.g Jimenez-Valverde et al., 2011). Other challenges arise from the sensitivity of SDMs to aspects such as study area size, the volume of data used to fit the models, and the spatial distribution of the data (aggregated/disaggregated) (e.g Phillips et al., 2007; Peterson et al., 2011; Guisan et al., 2017).

Species-specific responses to environmental gradients and spatial characteristics limit the automation and scalability of SDMs. Here we presented the AutoMaxEnt routine, a collection of tools and routines that allows for a flexible and fast exploration of multiple spatial features and scenarios as well as model adjustment, evaluation and selection metrics. The AutoMaxEnt routine uses the MaxEnt algorithm (Phillips et al., 2006) as its core element for the construction of species niche and distribution models.

### The AutoMaxEnt function

The AutoMaxEnt function is the main piece of this routine. This function is a big wrapper around the MaxEnt *dismo* R-package (Hijmans et al., 2023) with additional functions attached to deal with data formatting, model fit parametrisation, model evaluation, and selection. Therefore, the MaxEnt (Phillips et al. 2006) algorithm is at the core of this function. MaxEnt, for maximum entropy, is a presence-only species distribution modelling algorithm (Phillips et al., 2006). This algorithm has been widely used in species distribution modelling approaches where absence information is not available. Instead, to calculate the probability of occurrence of a species in a given area, MaxEnt draws a set of random background points from the environment, which are then compared with the presence data (Phillips et al. 2006).

Compared with other SDMs algorithms, MaxEnt presents a series of advantages that make it ideal for multi-species studies: (1) It is a semi-unsupervised machine learning algorithm with multiple adjustment and fit options; (2) Is a presence-only method, circumventing the scarcity of absence data (e.g Guillera-Arroita et al., 2015). This allows for a wider adoption of available species spatial records used for macroecological studies; (3) is a method that has been extensively used in ecology and conservation, and has been proven to produce more accurate results than other SDM methods with less data (Fitzgibbon et al. 2022); (4) it can work with highly multidimensional data; (5) it offers access to base parametrization and optimization, which make it flexible and customizable (Phillips et al. 2006). All this makes it ideal for automation.

### The AutoMaxEnt workflow

For the execution of AutoMaxEnt we just need species presence data in the form of an *sf* spatial object (Pebesma & Bivand, 2023) or a GBIF records file (https://www.gbif.org/), and spatial environmental data in the form of a raster (*raster* R package) or spatraster (*terra* R-package) object (Hijmans, 2024). Once this data is gathered and cleaned, AutoMaxEnt offers a series of options to further process the information. By default, the function will define the study area, which is the spatial region we want to use to define our species niche, as the bounding box around the minimum complex polygon defined by our distribution data (e.g Jarnevich et al., 2017) (Figure 1). However, we can also use a custom polygon to define this area (Figure 1) or add a buffer around any defined study area (Figure 1). The study area is going to contain all the information used for the model; the environmental information will be adapted to this study area, if needed.

**Figure 1.**
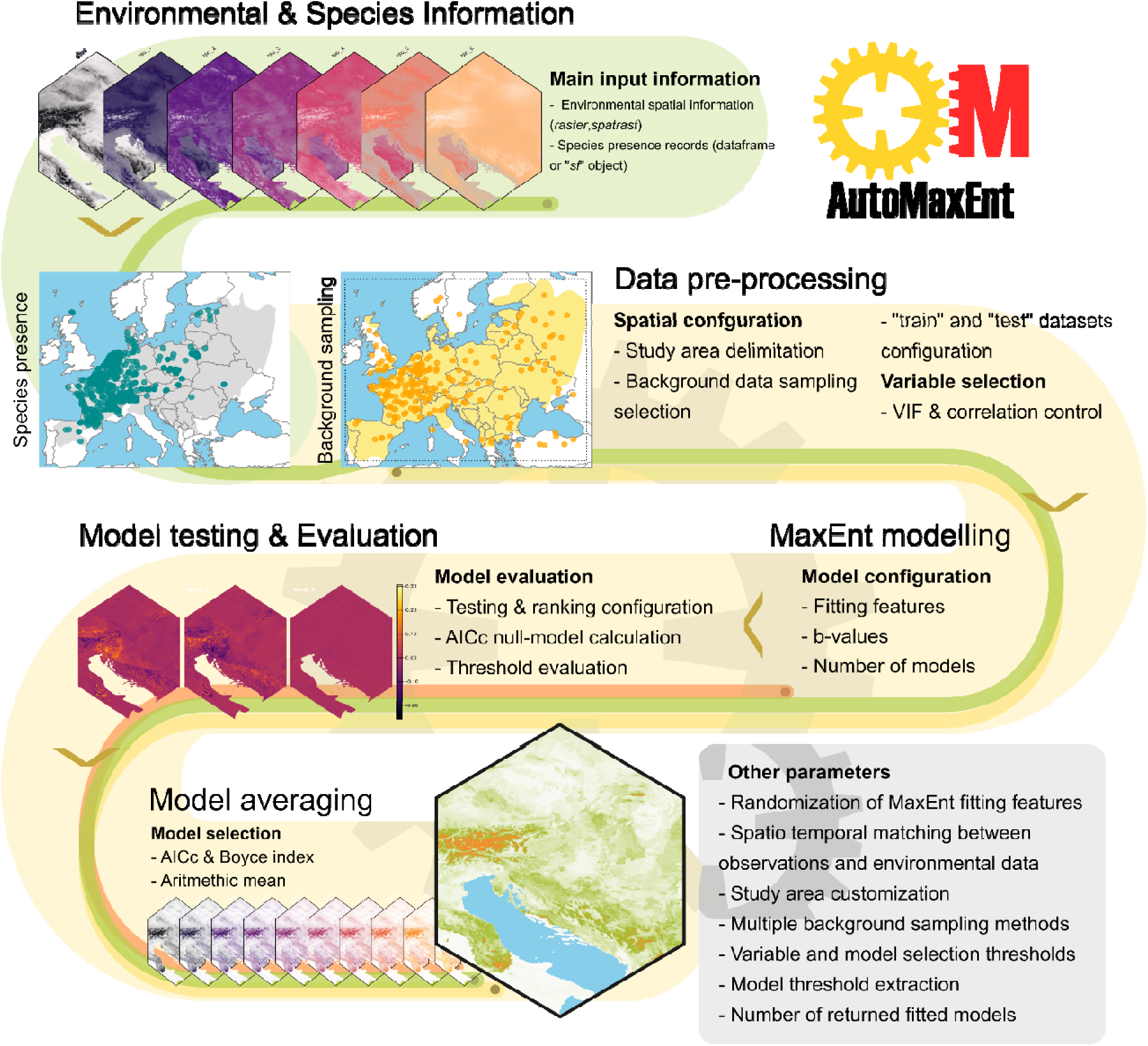
Main workflow structure of AutoMaxEnt. With little intervention from the user, AutoMaxEnt is able to get environmental and species distribution information, process this information and generate a wide diversity of Maximum Entropy models from which it will select the best performing ones to generate a final average of the likely distribution of the target species. Parameters regarding data processing, model fit and selection can be accessed and changed by the user, providing extra flexibility to explore different modelling scenarios.

Once the study region is defined, we need to sample this area to generate our background data. In this case, AutoMaxEnt offers four different sampling techniques:

1. Random sampling: points are randomly sampled within the study area. This is the default method used by MaxEnt (Phillips et al., 2006) (Figure S2.a).
2. Presence density sampling: points are sampled using a density matrix derived from the presence data (e.g Barbet-Massin et al., 2012; Jarnevich et al., 2017; Barber et al., 2022). This method favours the areas adjacent to species presences over areas in which the species hasn’t been recorded (Figure S2.b-S3.a).
3. Inverse density sampling: background points are drawn using the inverse of the density matrix derived from the species presences. Therefore, background points are more likely to be sampled away from the species presence data (Similar to Barbet-Massin et al., 2012) (Figure S2.c-S3.b).
4. Environmental sampling: in this case, background points are sampled using a density matrix derived from the first two principal components calculated using the set of environmental variables (similar to Da Re et al., 2023 and Broussin et al., 2024). In this case, sampling will be guided by the aggregation of environmental data, with the areas that present more common environmental conditions being sampled at a higher frequency than marginal or rare habitats (Figure S2.d-S3c-d).

With our presence and background data ready, the environmental information is extracted from the spatial objects. Different combinations of spatial data are likely to contain highly correlated variables, which can lead to collinearity, redundancy, and other fitting problems for the model (Zuur et al., 2010). Due to this, we have the option within AutoMaxEnt to run a variable selection routine. This variable selection is based on the Variance Inflation Factor (VIF) and variable correlations (Montgomery et al., 2021). First, VIF values are calculated for all variables; those that present a value of VIF above a threshold of 5 are discarded (this is a rule of thumb value widely used for this metric, e.g O’ Brien, 2007), and VIF values are recalculated until all variables present have a value below this threshold. The remaining variables are then tested for correlation using the Pearson coefficient (Galton, 1886). For any given pair of variables with correlation values greater than the set value (0.7 by default), one is removed at random to maintain all correlation values below our threshold.

Once our presence and background data are configured, our study area is defined, and the variables needed for the analysis are sampled and filtered AutoMaxEnt will proceed to fit different potential MaxEnt models. By default, MaxEnt uses a combination of multiple fitting features (linear, non-linear, quadratic, product, and hinge) to test and find the combination of parameters that better match our distribution and background data (Phillips et al. 2006). This produces highly reliable models and allows MaxEnt to fit complex data. However, it also limits the number of models fitted and can result in some overfitting (e.g Merow et al., 2013). Under the AutoMaxEnt routine, fitting functions are randomly suppressed, and a maximum of 4 are considered at once. Thus, instead of one, multiple MaxEnt models are fitted according to the different combinations of parameters (Figure 1). This produces a wider diversity of responses which, in theory, lead to a better representation of species distributions (e.g Redding et al. 2017; Albaladejo-Robles et al., 2025). Although this process is automatic within AutoMaxEnt, there are a few parameters and decisions that can be set by the user:

a. Number of models to fit (“n” hereafter): The number of models the user wants to produce. If the parameter “sampling” is not random, this field is overwritten (see below).
b. β-multiplier: this is a tuning parameter that controls feature expansion and regularisation strength in the model (Philips and Dudik, 2008; Merow et al., 2013). This parameter scales the regularisation penalty applied to each fitted class. Higher β-values prevent overfitting by penalising overly complex models, thus producing simpler or smoother models. Lower β-values impose weaker regularisation, allowing more complex models to form, potentially leading to overfitting. The β-parameter (or multiplier) is often used to balance model complexity and predictive performance (Merow et al., 2013). In AutoMaxEnt, the user can choose to fit a fixed β-value for all models or a range of parameters to test. Depending on the type of parameter sampling, these values (if there is more than one) will be sampled at random or systematically (see below).
c. Type of parameter sampling: AutoMaxEnt offers two ways to sample the space of fitting parameters: at random (the default), in which each model will contain a random set of fitting features and β-parameters, or based on combinations of fitting features and β-parameters. In the first scenario, several models, defined by “n”, with random features are created. The second option will create models with all the possible combinations of fitting parameters (form combinations of 1 up to 4) and β-parameters (in the case we have more than one specified). As a result, the “n”field is suppressed, and all the possible existing models are fitted.

With these parameters set, AutoMaxEnt will fit and evaluate the selected number of models, or the number of models resulting from all possible combinations of feature classes and β-parameters. By default, the function evaluates the models to generate comparison tables. Parameters such as the Area Under the Curve (AUC) or various cut-off values for the models, like Kappa (Kraemer, 2014) or the true statistical skill TSS (Allouche et al., 2006) are calculated and returned.

Most of these metrics are calculated internally by MaxEnt (Phillips et al., 2006) using different probability thresholds to build the initial confusion matrix. In addition to these metrics, AutoMaxEnt also calculates the Corrected Akaike Information Criteria (AICc, e.g Stoica and Selen, 2004) and the Boyce index (Boyce et al., 2002). The AICc measures model overall performance and fit. This metric is frequently used to compare models based on their goodness of fit and complexity (Cavanaugh and Neath, 2019). The AICc adds a penalisation parameter, favouring simpler and more generalised models over overly complex and overfitted ones. Within the AutoMaxEnt routine, the AICc is used to rank our models from best, lower AICc values to worst fit, higher values of AICc. For this, first, a null model is created with a randomly distributed sample of pseudo-presence and background data. This null model is then compared to the models fitted with real data and ranked amongst the others using the AICc. Models with an AICc higher than the null-model are known to be worse than a random fit and, therefore, can be later discarded if necessary.

To test model accuracy at classifying species presence and absence/background records, we used the continuous Boyce index (Boyce et al., 2002). This index has been proposed as a more reliable alternative than the AUC for binomial classifier models (e,g Liu et al., 2025; see also Lobo et al., 2008). The Boice index measures the correlation between predicted suitability and observed presence frequency data (Boyce et al., 2002). Since the metric is based on correlations, it varies from -1 to 1. A positive index value indicates that presence records are more frequent than expected by change in areas with higher predictive values, whereas values close to 0 mean that predictions are not better than random (Boyce et al., 2002). Negative Boyce index values indicate models that perform worse than random (Boyce et al., 2002). The Boyce index is threshold-independent and particularly suited for presence-only models such as MaxEnt (e.g., Liu et al., 2025).

If the user chooses to have a selection of “n” models returned by the function, instead of the total number of models, both the AICc and the Boice index are used to select the models (Knowalik et al., 2021; Stoice and Selen, 2004; Boyce et al., 2002). First, models are classified according to their AICc values from best to worst fit (lowest AICc to highest AICc values, respectively). Once all models have been ranked, the first “n” models are selected. After this first selection, models are filtered using the Boice index. By default, a threshold value of 0.5 is applied, which means that only models with an index above 0.5 are selected and returned by the function. This threshold can be set by the user and can have an impact on the number of models selected and the overall results. Because of this filtering, and even though “n” is fixed by the user, the number of final models can be lower. Similarly, if the null model is among the selected ones, a warning message is issued, and this null model is dropped from the selection.

For model testing and evaluation, a fraction of the presence and background data is extracted from the full dataset before model fitting. These two subsets of data comprise our test and training data. By default, 30% of data is put aside for testing purposes; the user can specify which percentage of data is used for training and testing. Testing data is used to internally evaluate the MaxEnt model accuracy and to calculate the Boyce index.

### Model averaging and function results

Once the models are fitted and the selection process is done, these are returned to the user in the form of an R “*list”* object. In addition to the MaxEnt models, all the information regarding the models’ parameters, their performance, and predictions is returned. Furthermore, the predictions of these models are averaged, using the arithmetic mean, and returned to the user, along with all the processed spatial information used to train and test the fitted models (Figure 2-3 & Table 1).

**Figure 2.**
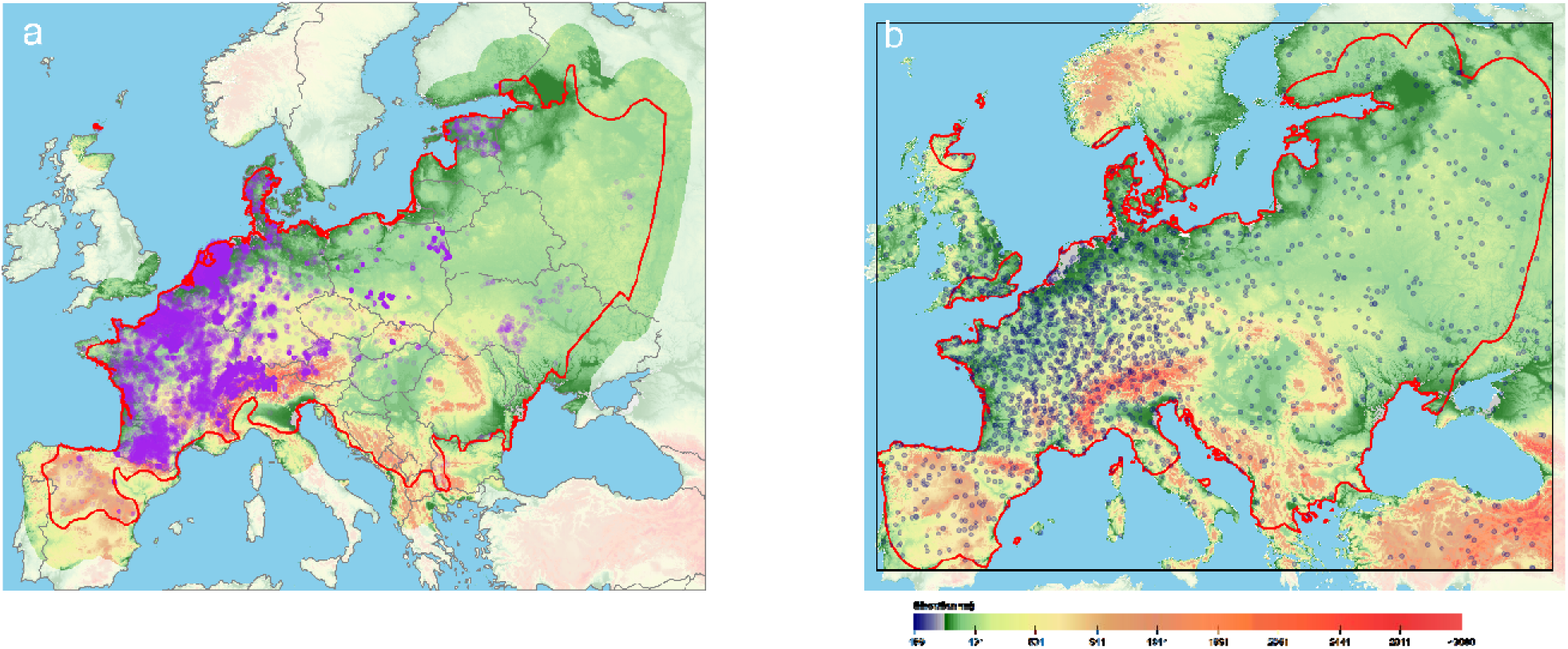
Distribution data for the common vole (*Microtus arvalis*). A) Georeferenced presence record (purple points) were downloaded from the Global Biodiversity Infrastructure Facility (GBIF) for the period 2000-2025 and later filtered to remove erroneous data (see Supplementary materials for details). The area highlighted in the map represents a buffer of 2 degrees around the range of the species (red line). This area will be used to define the complete study area or sampling region (black line in panel B). B) Distribution of background records across the study area (black polygon). Background points (dark blue points) were sampled using a spatial density kernel based on the spatial aggregation of presence data (Barbet-Massin et al., 2012; Jarnevich et al., 2017; Barber et al., 2022). This information is overlaid over a digital elevation map of ∼4.6 km pixel resolution (Fick and Hijmans, 2017).

**Figure 3.**
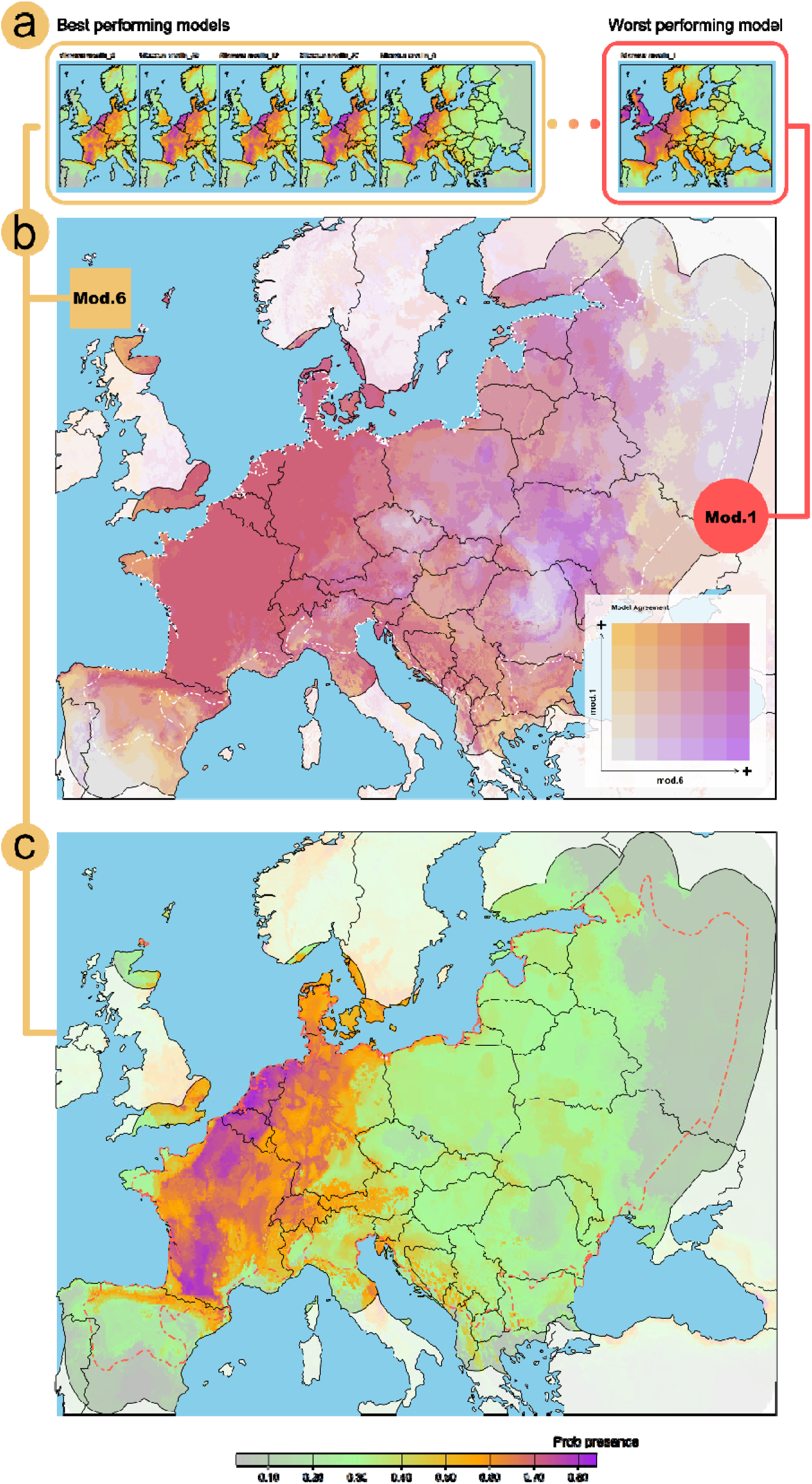
Potential distribution of M. arvalis in mainland Europe according to the AutoMaxEnt. Panel a shows the five best-fitted and performing models (gold) alongside the worst-performing model (red) (Table 1 & S3). Congruence between the “best” and “worst” models is shown in the bivariate map shown in panel **b**. Grey and red tones represent areas of high congruence between the models, for low to high probability of species occurrence (diagonal line in the **model agreement legend**). Purple and golden tones represent a higher probability of occurrence for model 6, the “best” model, and model 1, the “worst” one, respectively. Panel **c** shows the final model, which results from averaging the 5 top-performance MaxEnt models. In all cases, black solid lines represent the political borders, whereas dashed white lines, in panel b, and red, in panel c, represent the species range as presented in the IUCN Red List of Threatened Species (iucn.org). In panels **a** and **b**, only the results within the study area (species range map + 2 degrees) are highlighted, whereas other areas are partially masked.

**Table 1.** Performance and fit metrics for the 5 best-performing models (**id**). Models are ranked according to their corrected Akaike information criteria (**AICc**). Differences between all models and the one with the lower AICc value are also returned (Δ **AICc**) along with other parameters such as number of model coefficients (n-coefs) and log-odds (Log Odds). Model accuracy is internally evaluated by MaxEnt using the Area Under the Curve (AUC), both for the train and test datasets (**Train AUC** and **Test AUC,** respectively). The differences between the two AUC metrics represent the level of overfitting of the model. MaxEnt also computes the True Statistical Skill threshold (**TSS MaxEnt**). In addition, independent tests are used within AutoMaxEnt to re-calculate the TSS threshold (**TSS threshold**) and extract the TSS mean (**TSS mean**) and max values (**TSS Max**). Model classification performance is evaluated using the continuous Boyce index (**Boyce**) implemented within the AutoMaxEnt routine. AutoMaxEnt also returns the list of MaxEnt parameters used to fit each model (e.g the list of fitting features, b-values, etc). The full list of model parameters can be consulted in Table S3.

| mod.i<br>d | Train<br>AUC | Test<br>AUC | Diff<br>AUC | TSS<br>MaxEnt | TSS<br>threshold | TSS<br>mean | TSS<br>max | Boyce | n-coefs | Log Odds | AICc | Δ AICc |
| --- | --- | --- | --- | --- | --- | --- | --- | --- | --- | --- | --- | --- |
| 6 | 0.67<br>0 | 0.66<br>2 | 0.00<br>9 | 0.565 | 0.540 | 0.24<br>3 | 0.42<br>8 | 0.999 | 149.00<br>0 | -<br>65914.78<br>4 | 132133.62<br>3 | 0.000 |
| 39 | 0.66<br>3 | 0.65<br>8 | 0.00<br>5 | 0.596 | 0.580 | 0.23<br>2 | 0.41<br>9 | 0.998 | 79.000 | -<br>66302.18<br>9 | 132764.07<br>3 | 630.451 |
| 12 | 0.66<br>3 | 0.65<br>8 | 0.00<br>4 | 0.569 | 0.570 | 0.23<br>6 | 0.41<br>3 | 0.999 | 95.000 | -<br>66297.10<br>6 | 132786.66<br>5 | 653.042 |
| 27 | 0.65<br>9 | 0.65<br>7 | 0.00<br>2 | 0.609 | 0.590 | 0.23<br>0 | 0.41<br>0 | 0.998 | 58.000 | -<br>66511.01<br>6 | 133138.94<br>7 | 1005.32<br>5 |
| 8 | 0.65<br>8 | 0.65<br>5 | 0.00<br>2 | 0.573 | 0.560 | 0.22<br>4 | 0.39<br>8 | 0.998 | 62.000 | -<br>66526.74<br>1 | 133178.52<br>8 | 1044.90<br>5 |

### Workflow example

To illustrate the overall functionality of the AutoMaxEnt function, we are going to build an SDM of the Common vole (*Microtus arvalis*). This rodent species is widely distributed across Europe and certain regions of Asia and presents a fair number of records in the Global Biodiversity Infrastructure Facility (GBIF, https://www.gbif.org/) as well as spatial range information in the IUCN Red List of threatened species (https://www.iucnredlist.org/), where the species has a status of least concern (Figure 2.a).

To model the distribution of this species, we extracted more than 34,000 presence records from GBIF for the period 2000-2025. These original records were then cleaned and selected to assure maximum verisimilitude of the presence data (see supplementary materials for a detailed description). As a result, we obtained 10,771 records for the analysis. Despite their commonality, *M. arvalis* presence data are highly clustered across central Europe, and it is scarce elsewhere in the species’ range (Figure 2).

To describe the habitat of *M. arvalis,* we downloaded climatic information from BIOCLIM (version 2.1) (Fick and Hijmans, 2017). BIOCLIM is part of the WordClim dataset and contains 19 different climatic variables of biological interest (see supplementary materials for a full list). These variables are calculated for the whole world at a resolution of 1km (Version 2.1). In addition, we also got elevation data from the SRTM 90m DEM Digital Elevation Database (NASA JPL, 2013). This elevation data is included within WorldClim and is the result of the aggregation of data from the Shuttle Radar Topography Mission (SRTM) (NASA JPL, 2013) and the GTOPO30 (USG,1996) (Fick and Hijmans, 2017). We obtained all the environmental information at a resolution of 2.5 arc min (∼4.6 km at the equator) using the *geodata* R-package version 0.6-2 (Hijmans et al.,2024).

### Setting up the Function parameters

AutoMaxEnt works mostly automatically once the spatial information (species records and range) and the environmental data (explanatory variables) are selected. However, it is always useful to tweak the function parameters based on the information we have about the species and the data we are using. In this case, we know that *M. arvalis* is widely distributed across its entire range and that the patterns of observations we are getting are probably due to some sampling bias (Figure 2.a).

Due to this aggregation of observations, we selected an observation-driven background sampling to produce the background data (“*BwData”* in AutoMaxEnt). The number and distribution of background information can have a profound impact on model performance and transferability (e.g Whitford 2024; Steen et al., 2024). By choosing a density-driven background sampling, we intend to control observation bias while extracting a significant amount of environmental information. We set the number of background points to 10,000, which is close to a 1:1 ratio with the presence data and is also the default value for the function (and for MaxEnt) (Figure 2.b).

Since we have range information for the species, this will be used to delimit the study area (Figure 2). This is the region from which both the environmental and background information will be extracted (Figure 2.b). For this experiment, we want our model to be able to produce predictions for the future. With climate change producing latitudinal shifts in climatic gradients, we want to add a few degrees of extra area to our study area. This will ensure that we capture larger climatic variation in our background data, and therefore, in our model. For this example, we are going to configure AutoMaxEnt to add a buffer of 2 degrees around our study area (∼222 km at the equator) (Figure 2.b).

Although BIOCLIM includes climatic variables of biological interest, a large portion of them will exhibit higher levels of correlation and collinearity (Supplementary materials Table S1 and S2). This is likely to add redundancy to our models and will eventually lead to overly complex or overfitted models, among other potential problems (Zuur et al., 2010). To reduce the dimensionality of our predictors, we are going to set AutoMaxEnt to perform a variable selection using the default parameters for VIF (5) and correlation (0.7) (e.g Patiño et al., 2023; Albaladejo-Robles et al., 2025).

For the rest of the analysis, we will select a range of β-multiplier values between 1-5 and run a total of 50 models with a random combination of model-fitting features (Table 1, full results shown on Table S3). By setting a low range of β-multiplier values, we ensure that complex or overfitted models can be formed through the MaxEnt fitting. Since *M. arvalis* presents a large distribution area and observations are clearly biased, limiting the regularisation and feature expansion penalisation is a way to narrow down the model predictions. Since we want to retain the parameters and the model performance metrics for all models, we are not going to run an automatic model selection. Instead, once calculated, we will select the 5 best-performing models manually (using the same parameters as the default AutoMaxEnt variable selection parameters) to calculate the average. To do this, we set the model selection feature of AutoMaxEnt off and fix a Boice index value of 0.7 for the selection.

## Results and model averaging

With the present configuration, AutoMaxEnt has returned 29 unique models with varied performance, internal configuration, and predictive power (Table 1 & Table S3). This is lower than the initial objective (50) because, due to the random nature of the parameter selection and the low number of β-values, some parameters combinations were duplicated and produced identical models. Those duplicated models were discarded from the final set (Table S3). Overall, models present high levels of predictive power, with Boyce index values ranging from 0.99 to 0.69, with the worst-fitted model presenting an AICc 2.54% higher than the model with the best fit (Table S3). The best 5 performing models (Table 1 & Figure 3.a) showed very high Boyce index values, all within the range of 0.99, and an absolute difference in AICc between the “best” and “worst” fitted model of 0.79% (Table 1 & S3). Despite these higher values, differences between MaxEnt train and test AUC values were low (mean difference of 0.005 ± 0.002), suggesting no model overfitting (Table 1). These high Boyce index values can raise some concerns, since they indicate an almost perfect predictive performance of the models. Although suspicious, these values match the observed TSS threshold values returned by the confusion matrix (Table 1 and S3). All models present TSS cut-off values in the range of 0.49 to 0.6. A lower probability threshold indicates a weak environmental filtering of the species (Allouche et al., 2006). This low environmental response allows for a wider area of presence to be estimated (Figure 3.c), which then allows for higher levels of sensitivity (correctly predicting presence areas) but creates a decrease in specificity (erroneous predictions of background locations) (Allouche et al., 2006). This can be observed in the TSS maximum values for all models that ranged from 0.33 to 0.42 (Table 1 & S3) and in the relatively low performance exhibited by MaxEnt’s AUC calculations (Table 1 & S3). This helps explain the high values of the Boyce index, provides an approximation to the role of the selected environmental variables on the distribution of the species, and shows why it is important to rely on multiple metrics to evaluate model performance (e.g Knowalik et al., 2021; Stoice and Selen, 2004).

Those performance results were expected, due to the generalist character of the species and the abundance of data, although clearly clustered, within its range (Figure 2.a). Consequently, model predictions showed broad distributions, making most areas around central Europe highly suitable for the species (Figure 3.a-c). This pattern was observed across all 33 models (Figure S4) with different degrees of intensity and granularity (the intensity in the change of probability of occurrence between adjacent pixels). The degree of agreement between the “best” and “worst” models was also high (Figure 3.b), despite the difference in granularity between them (Figure 3.a-b). The best performing models models were combined and averaged to produce the final prediction for the potential distribution of *M. arvalis* in Europe (Figure 3.c).

The final combined distribution of *M. arvalis* shows a clear distribution pattern towards the central and northern regions of Europe, with high probabilities of occurrence registered as north as the Orkney Islands (Figure 3.c). This matches the general spatial information of the species, with relatively abundant populations in Northern Spain, France, Germany, the Netherlands, Belgium, Switzerland, and Denmark (Amori, 2024). However, the distribution seems to be truncated towards the eastern European region, although this matches well with the IUCN spatial information (Amori, 2024), it underrepresents the distribution of the species in this area (Stakheev et al., 2023). However, this can be explained by the omission of many records for Ukraine and Russia (Figure 2.a) (Stakheev et al., 2023), and the extent of the IUCN range map for the species, which seems to underrepresent the area in which the species is found across the black sea and western Russia (Amori, 2024; Stakheev et al., 2023). This is likely due to the initial filtering process of presence records (see supplementary materials S1), and for some studies or collections being absent in GBIF.

Generally, AutoMaxEnt has successfully reconstructed the potential distribution of *M. arvalis*, using a multi-model averaging approach. We’ve been able to produce high-performing models using a simple set of function parameters and minimum user effort in terms of data preparation, model adjustment, evaluation and selection. Although results follow the known information for the species, some limitations, common to any SDM model (Peterson et al., 2011), arose as a function of the quality of the species spatial information and the selection of variables, which has been done arbitrarily.

## Conclusions

AutoMaxEnt presents an easy and flexible routine to build and explore species distribution models using MaxEnt. The built-in tools for data integration, preparation, and evaluation allow for a cohesive integration of study area definition, variable selection and data preparation for the analysis. By integrating these functions with different background sampling techniques and multiple parameters to define the spatial domain of the analysis, AutoMaxEnt allow users to explore multiple study areas and sampling scenarios efficiently by changing just a few function parameters. Furthermore, AutoMaxEnt integrates this data pre-processing pipeline with environmental variable selection, model fitting, evaluation and model selection. Thus, creating a cohesive workflow that allows users with different levels of technical expertise to produce reproducible, high-quality species distribution scenarios. This makes AutoMaxEnt particularly valuable for large-scale, multi-species or multi-scenario species studies.

A key advantage of AutoMaxEnt lies in its flexibility. Although the routine is designed to run automatically with sensible default settings, every major modelling decision from study area construction to background sampling, feature class selection, β-multipliers, and model filtering can be easily modified by the user. This flexibility is directly built into the structure of AutoMaxEnt, which, rather than being a closed ecosystem, is formed by multiple functions wrapped around the *dismo* MaxEnt implementation (Hijmans et al., 2023). Therefore, the collection of functions that form this pipeline can be used in isolation to fit a different SDM algorithm, process spatial information, evaluate model performance, create background or pseudo-absence data, etc. Thus, allowing users to create other SDM or analytical pipelines using these tools. This also allows users to access further parameters that are set by default in AutoMaxEnt. For example, density kernels for the sampling of background data are set by default to use a 1000 by 1000 matrix and a bandwidth of 10% the extent of the study area (see Figure S3). Additionally, AutoMaxEnt is easily scalable to high computer performance platforms, allowing for the adjustment of hundreds of SDMs simultaneously.

This balance between automation and customisability empowers researchers to tailor the modelling pipeline to their needs or the requirements of specific hypotheses. Such flexibility is essential for addressing species-specific responses to environmental gradients and ensuring that automated workflows do not sacrifice ecological realism for convenience. Due to this, we believe that AutoMaxEnt position it as a powerful collection of tools for modern species distribution modelling. By reducing the need for manual preprocessing while retaining full analytical control, the routine facilitates scalable, reproducible, and ecologically meaningful modelling across broad taxonomic and geographic contexts. As biodiversity assessments increasingly require automated yet robust workflows, AutoMaxEnt contributes a practical and adaptable solution for large-scale SDM research and its applications in ecology and conservation.

## Notes

https://github.com/BioDivHealth/AutoMaxent/tree/main

### Competing Interest Statement

The authors have declared no competing interest.

