## Supplementary materials for "Towards the automation of Species Distribution Modelling: The AutoMaxEnt routine"

**1 GBIF records filtering and correction**

The Global Biodiversity Infrastructure Facility (GBIF) is formed from the aggregation and standardization of multiple datasets and data collection platforms. GBIF contains species information ranging from individual scientific studies to government-driven census, citizen science, private collection and museum records. This wide diversity of sources makes GBIF the reference platform for the collection and the deposit of species records information, but it also adds a lot of uncertainty to its data. Due to this, many spatial records are prone to contain inaccurate or erroneous information (Beck et al., 2014).

GBIF inaccuracies can take many forms: rounded coordinates or coordinates derived from regions or protected areas centroids; erroneous or mismatched metadata; large spatial inaccuracies; erroneous or missing times of recording; and many other factors can impact the quality and veracity of the records collected by this platform (e.g Beck et al., 2014; Zizka et al., 2019). Considering the sensitivity of any species distribution model (SDM) to input data, an intense filtering and selection of the data is needed before conducting any analysis. To preserve only the more accurate records for *Microtus arvalis,* we performed a multi-step data filtering procedure. First, we use the R-package *CoordinateCleaner* version 2.0-20 (Zizka et al., 2019) to remove the observations with conflicted or erroneous coordinates, metadata mismatches, and invalid locations (museum and private collections, zoos, etc). After this first filtering, we used the IUCN Red List of Threatened Species to filter out the points recorded outside the species range. Even though IUCN range maps don’t reflect the real distribution of the species, by using them for the filtering of species records, we can, indirectly, add some expert-based information into this process.

Additionally, AutoMaxent also impose a series of internal filtering processes, which can be in part controlled by the user. First, AutoMaxent includes the option to remove or preserve the duplicated spatial records. In many cases, spatial records can be tied to protected areas, centroids, sampling stations, or regional spatial centroids. This aggregation can create some spatial bias that we might want to remove. In those cases, we can decide to remove these spatially redundant records. This is the case with the example case study of *M. arvalis*. Another case in which AutoMaxent filters information is when the time matching option is implemented. In this case, records with wrongly formatted dates are discarded from the analysis. In addition to this, some background sampling methods, when used with time-matching, also impose a strict selection on the number of records used. For example, if we run time-matching with an observation density-driven background sampling, if we have less than 3 spatial records (the minimum to compute a density kernel) for any given time, those observations and the period are ignored. This illustrates how data quality and model parameter requirements shape data filtering.

**2 Environmental variables**

Variable selection has a huge impact on species distribution modelling and niche reconstruction studies (e.g Austin et al., 2011; Petitpierre et al., 2017; Diaz-Vallejo et al., 2024). Therefore, the way in which we select these variables should reflect both the capabilities of our model to handle the data and the ecological needs of the species (Mancini et al., 2025). Ideally, we want to include as much relevant information for the species as possible. This can take the shape of climatic, land-cover, dispersal and biological spatial information. For the example *M. arvensis* we have used a combination of climatic and terrain information in the form of BIOCLIM (Karger et al. [2017](https://onlinelibrary.wiley.com/doi/10.1002/ece3.71580#ece371580-bib-0016)) and elevation data (Fick and Hijmans, 2017).

For the modelling of the potential distribution of *M. arvensis*, we selected a combination of climatic and topographic variables. For the climatic variables, we used the Bioclim dataset (Karger et al. [2017](https://onlinelibrary.wiley.com/doi/10.1002/ece3.71580#ece371580-bib-0016)) (<https://chelsa-climate.org/bioclim/>). BIOCLIM climatic time series spans from 1990 to 2021 (version 2.1) (Fick and Hijmans, 2017). These time series contain yearly values of seasonality and annual trends, as well as extreme or limiting factors such as minimum and maximum temperatures, calculated from monthly average values (Fick and Hijmans, 2017). BIOCLIM contains 19 different climatic variables with ecological potential; Annual mean temperature (Bio1); Mean diurnal range (Bio2); Isothermality (Bio3); Temperature seasonality (Bio4); Max temperature of the warmest month (Bio5); Min temperature of coldest month (Bio6);  Temperature annual range (Bio7); Mean temperature of wettest quarter (Bio8); Mean temperature of direst quarter (Bio9); Mean temperature of warmest quarter (Bio10); Mean temperature of coldest quarter (Bio11); Annual precipitation (Bio12); Precipitation of wettest month (Bio13); Precipitation of driest month (Bio14); Precipitation Seasonality (Bio15); Precipitation of wettest quarter (Bio16); Precipitation of driest quarter (Bio17); Precipitation of warmest quarter (Bio18); and Precipitation of coldest quarter (Bio19).

To complement this data, we also used the elevation data from WorldClim (worldclim.org). This elevation data is derived from a digital elevation model (DEM) created by the Shuttle Radar Topography Mission (SRTM) (NASA JPL, 2013) and the GTOPO30 (USG,1996) (see Fick and Hijmans, 2017 for more details). All spatial information had a global coverage and was transformed to the same grid resolution (~4.6 km at the equator) and cropped to the extent of our study area by AutoMaxent (see Figure S1.F or Figure 2 in the main text for more details).

Although MaxEnt can deal with highly dimensional data, most of our climatic and topographical variables are going to present a high degree of correlation between them. This can lead to model inaccuracies and overfitting (e.g Zuur et al., 2010). Additionally, since we are not doing a methodological review of the factors influencing the species' presence, we are going to reduce the dimensionality of our explanatory variables using the statistical methods implemented in AutoMaxent. First, a variation inflation factor (VIF) sequential analysis is implemented. The VIF is a metric that measures the variable collinearity. As a rule of thumb, VIF values should be below 5 to be considered as having low collinearity (O’ Brien, 2007). This is the threshold AutoMaxent uses as its default to include or exclude the variables. In a first selection step, AutoMaxent calculates the VIF for all the explanatory variables loaded into the function. Variables with a VIF value above the threshold are excluded, and the VIF is recalculated for the remaining variables. This process is repeated until all variables have a VIF value of 5 or below (Table S1). After this first selection of variables is finished, AutoMaxent calculates the correlation between pairs of variables and excludes, at random, one of each pair of variables with a correlation value greater than 0.7 (e.g Patiño et al., 2023) (Table S2). An earlier version of this protocol can be found in Albaladejo-Robles et al (2025).

As a result of this filtering, 8 environmental variables were selected: Bio2, Bio4, Bio5, Bio9, Bio15, Bio18, Bio19, and elevation. All with VIF values below 5 (Table S1) and pairwise correlations lower than 0.7 (Table S2). This selection of variables has been numerically driven and, therefore, might not reflect the real ecological preferences/needs of the species. All variables were evaluated using the subset of locations belonging to the species records and background data. This is implemented in AutoMaxent to save computation time and to evaluate the data that is going to be used for the training and testing of the model. This is implemented in AutoMaxent to save computation time and to evaluate the data that is going to be used for the training and testing of the model.

**Table S1.** Variance inflation factor (VIF) matrix of the set of environmental variables considered for the modelling of the potential distribution of *M. arvensis* in its distribution area (Figure 2.a). Rows contain the individual VIF values for a given combination of environmental variables. Columns represent each iteration of the VIF variable reduction process. At each step, the variable with the highest VIF is removed until all variables present a VIF value below 5. Selected variables are marked in **bold**. A full transcription of the variable codes can be found in section S.2.

| **Variables** | **1** | **2** | **3** | **4** | **5** | **6** | **7** | **8** | **9** | **10** | **11** | **12** | **13** |
| --- | --- | --- | --- | --- | --- | --- | --- | --- | --- | --- | --- | --- | --- |
| Bio1 | 703.7 | 750.8 | 647.0 | 98.2 | 68.4 | 68.0 | 7.8 | 7.6 | 8.0 | 7.3 | 7.2 | 6.5 |  |
| Bio10 | 1352.1 | 1278.8 | 750.8 |  |  |  |  |  |  |  |  |  |  |
| Bio11 | 2033.6 | 2104.7 |  |  |  |  |  |  |  |  |  |  |  |
| Bio12 | 121.0 | 110.7 | 124.1 | 112.3 | 108.8 | 80.3 | 75.1 |  |  |  |  |  |  |
| Bio13 | 66.2 | 59.7 | 66.7 | 64.2 | 64.5 | 35.6 | 34.6 | 21.1 | 17.5 | 19.2 |  |  |  |
| Bio14 | 39.3 | 35.4 | 37.1 | 36.0 | 36.0 | 35.1 | 33.3 | 33.8 | 17.3 | 16.1 | 16.0 |  |  |
| **Bio15** | 13.5 | 14.3 | 13.5 | 13.4 | 12.4 | 11.3 | 11.2 | 11.5 | 7.9 | 7.9 | 4.6 | 1.5 | **1.5** |
| Bio16 | 127.3 | 114.9 | 133.4 | 124.5 | 123.4 |  |  |  |  |  |  |  |  |
| Bio17 | 73.3 | 69.6 | 70.5 | 71.2 | 67.7 | 61.2 | 60.2 | 57.3 |  |  |  |  |  |
| **Bio18** | 19.9 | 19.3 | 18.2 | 19.2 | 17.3 | 16.0 | 14.3 | 11.9 | 11.9 | 12.0 | 7.7 | 3.4 | **3.4** |
| **Bio19** | 26.5 | 24.7 | 27.1 | 26.9 | 24.6 | 24.5 | 22.5 | 14.9 | 13.9 | 13.5 | 7.7 | 4.3 | **4.3** |
| **Bio2** | 90.2 | 82.7 | 78.4 | 78.0 | 53.9 | 53.8 | 24.8 | 25.4 | 25.3 | 2.6 | 2.6 | 2.6 | **1.7** |
| Bio3 | 63.2 | 66.4 | 55.9 | 60.6 | 54.9 | 54.0 | 41.8 | 41.9 | 41.4 | NA | NA | NA | NA |
| **Bio4** | 1284.8 | 1322.9 | 231.2 | 95.7 | 35.8 | 35.5 | 36.6 | 35.9 | 35.9 | 3.3 | 3.7 | 3.4 | **2.6** |
| Bio5 | 5000 > | 154.1 | 161.2 | 109.8 | 86.7 | 86.3 | NA | NA | NA | NA | NA | NA | NA |
| Bio6 | 5000 > | NA | NA | NA | NA | NA | NA | NA | NA | NA | NA | NA | NA |
| Bio7 | 5000 > | 189.0 | 187.3 | 187.0 | NA | NA | NA | NA | NA | NA | NA | NA | NA |
| **Bio8** | 2.7 | 2.8 | 2.8 | 2.7 | 2.7 | 2.6 | 2.5 | 2.6 | 2.6 | 2.6 | 2.5 | 2.5 | **2.3** |
| **Bio9** | 5.7 | 5.8 | 5.3 | 5.3 | 5.2 | 5.1 | 5.3 | 5.2 | 5.4 | 5.1 | 5.2 | 5.4 | **4.4** |
| **elev** | 6.1 | 5.5 | 4.7 | 4.6 | 4.5 | 4.4 | 4.3 | 4.5 | 4.5 | 3.8 | 3.3 | 3.1 | **1.6** |

**Table S2.** Correlation matrix for the environmental variables considered (first selection) for the modelling of the potential distribution of *M. arvensis* in its distribution area (Figure 1). Only variables selected after the VIF analysis (**Bold** variables in Table S1) are included. A full transcription of the variable codes can be found in section S.2

|  | Bio2 | Bio4 | Bio8 | Bio9 | Bio15 | bio18 | bio19 |
| --- | --- | --- | --- | --- | --- | --- | --- |
| Bio4 | 0.23 |  |  |  |  |  |  |
| Bio8 | 0.24 | 0.50 |  |  |  |  |  |
| Bio9 | 0.28 | -0.55 | -0.35 |  |  |  |  |
| Bio15 | 0.06 | 0.48 | 0.31 | -0.19 |  |  |  |
| Bio18 | -0.17 | 0.19 | 0.10 | -0.50 | 0.10 |  |  |
| Bio19 | -0.14 | -0.48 | -0.59 | 0.42 | -0.27 | 0.32 |  |
| Elev | 0.09 | 0.06 | -0.24 | -0.11 | 0.10 | 0.45 | 0.33 |

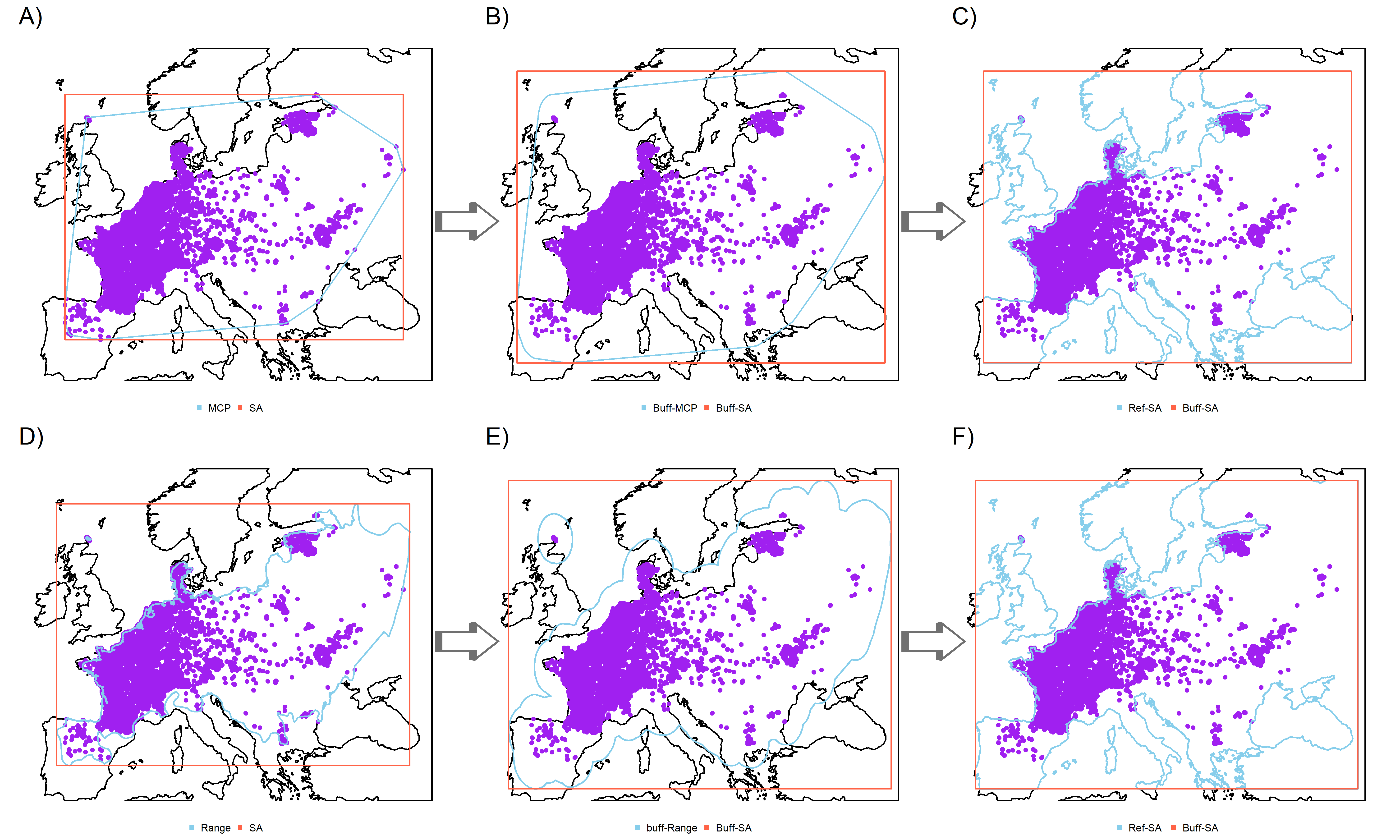

**Figure S1.** Workflow and reshaping of the study area to constrain environmental variables and delimit background sampling. The MaxEnt study area can be defined based on the minimum complex polygon defined by the species records (purple points) (A-C) or based on spatial information such as the IUCN Red List maps (https://www.iucnredlist.org/) (D-F). In all cases, the study area is defined by the bounding box (red polygon) formed around the spatial information (MCPs and range data in blue). In both cases, AutoMaxent allows the user to increase the study and sampling area by adding a buffer around the spatial information (B and E). Similarly, these study areas can be refined to avoid bodies of water or other limits that the users want to impose. Panels C and F show the final study regions once the spatial information is corrected by line coasts (blue lines). As can be seen, routes A-C and D-F produce different study areas and therefore distinct environmental variables, cutting points and background sampling results.

**3. Background data**

Although MaxEnt is considered a presence-only SDM method, the algorithm still needs to generate background information to compare the presence data against a sample of environmental points. By default, MaxEnt creates a set of 10,000 randomly distributed points across the study area. However, how these points are created and their distribution can impact the accuracy and performance of the final model (Barbet-Massin et al., 2012). This is particularly important when imperfect sampling occurs, or the presence records are highly aggregated (both cases present in our data). To evaluate the impact of the background data on our SDM models, as well as to select the best set of background points, AutoMaxEnt can create these points using four different methods across the distribution area of our target species (in this case, *M. arvalis*):

1. Background points were randomly distributed (Figure S2.a). This is the default approach of MaxEnt when creating the set of background points.

2. Background points based on *M. arvalis* presence points, where a density matrix is built based on the distribution of the species and background points are distributed according to this density matrix (Barbet-Massin et al., 2012; Jarnevich et al., 2017; Barber et al., 2022) (Figure S2. b).

3. Background points based on the inverse of a density matrix built based on the presence records of the target species (following the same principles as Barbet-Massin et al., 2012, when creating pseudo-absence data) (Figure S2. c).

4. Background points are distributed according to a density surface derived from the spatial aggregation of the first two principal components of a Principal Component Analysis applied on the environmental space covered by the study area (similar to Da Re et al., 2023 and Broussin et al., 2024) (Figure S2. d). As a result, the environmental space is sampled according to the general or dominant conditions of the environment. Therefore, the regions with more common or dominant environmental conditions are sampled, and the areas with rare or marginal environmental conditions are less frequently sampled (Figure S2. d). In this case, the background sampling combines elements from Da Re et al. (2023) and Broussin et al. (2024) in that it is not random environmental sampling and is not driven by species niche preferences.

In all cases, AutoMaxent use the study area provided, or internally created by the function, to delimit the background sampling area. This can be defined, expanded and subsequently refined using the different AutoMaxEnt parameters accessible by the user. These are described in more detail in the Tutorial (see link at the end of this document). Similarly, each background sampling method can be accessed through the “BackgroundPOINST” function. Thus, allowing the users to easily adapt these sampling methods for other presence-background or pseudo-absence SDM methods.

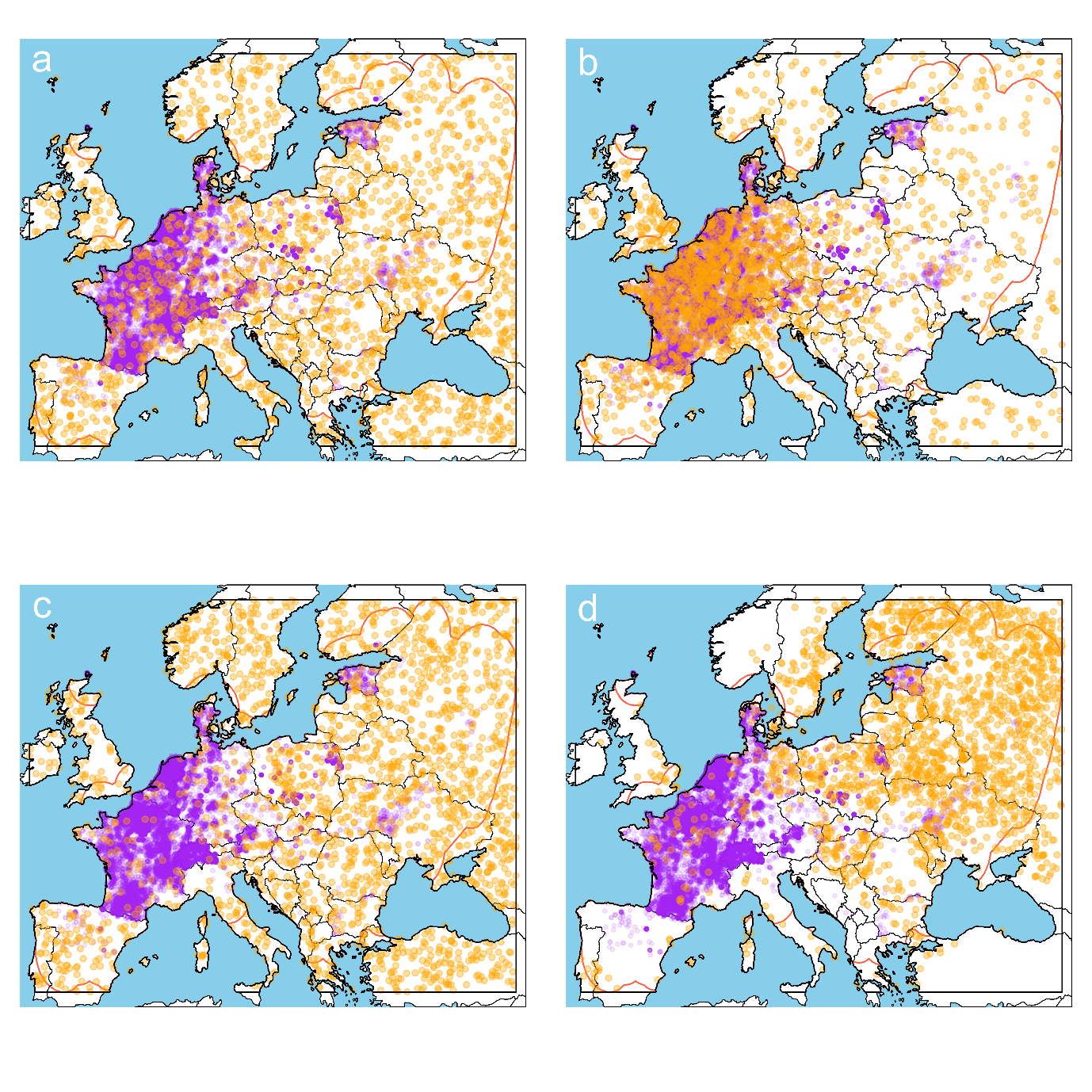
Figure S2. Distribution of *M. arvalis* presence records (purple points) and background information (orange points). The different panels show distinct background sampling strategies over the same sampling area (black box border); **a**, Random sampling of background locations across the study area; **b**, Background sampling based on a density matrix derived from the presence data of *M. arvalis*; **c**, sampling based on the inverse of the density matrix calculated in **b**; and **d**, a background site selection based on the environmental multidimensional space defined within the study area.

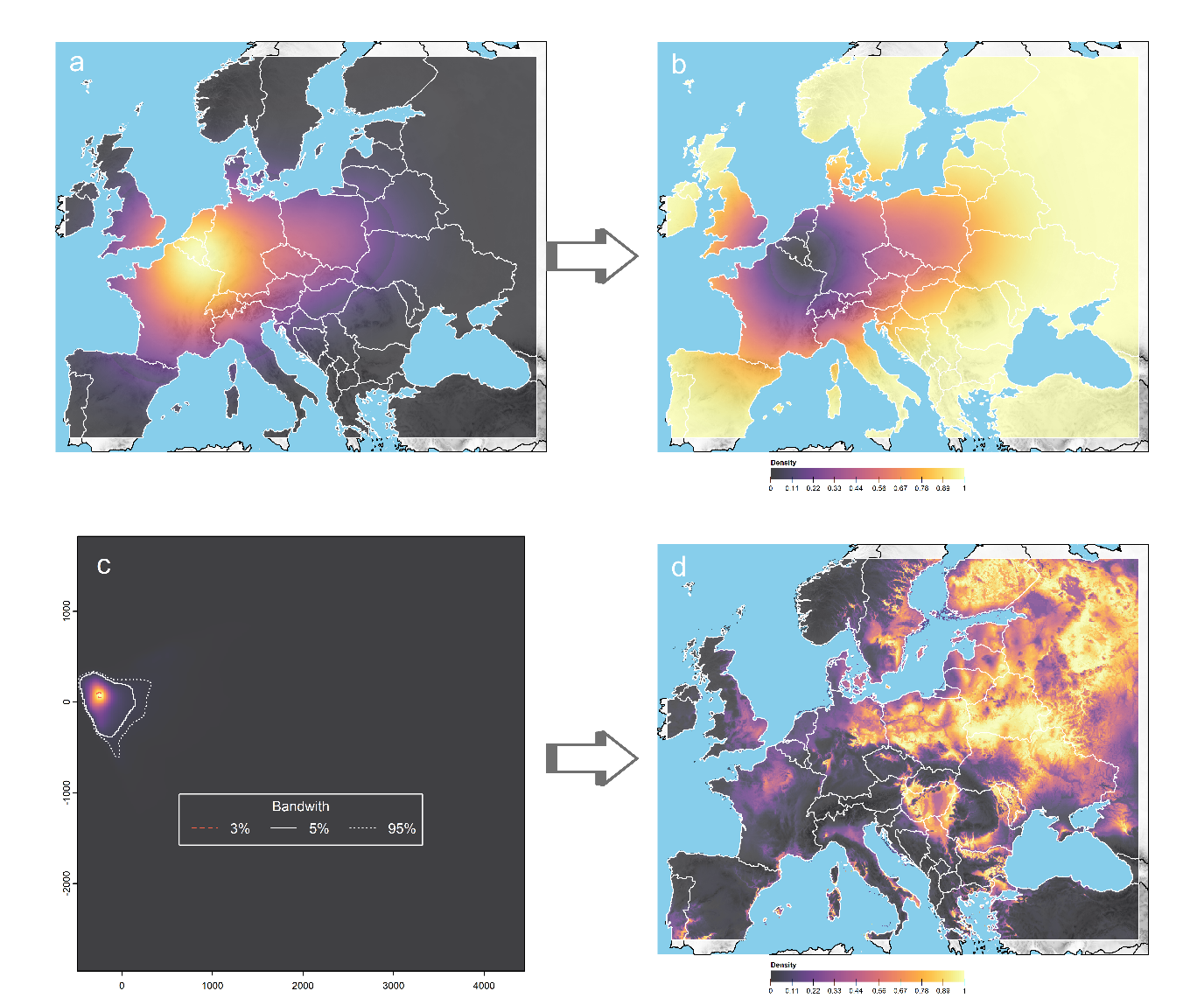

Figure S3. Density kernels were used for the background sampling of *M. arvensis* within the selected study area. Panel **a** presents a density kernel built using the presence data of *M. arvensis* within the selected study area. This corresponds to the “BwData” background sampling method in AutoMaxent (Figure S2.b). In panel **b**, the inverse of this density kernel is used to run the background sampling based on the inverse of the presence data background sampling (“BwData_inv” in AutoMaxent) (Figure S2.c). For the background sampling based on the environmental conditions, a PCA analysis is run, and score values of the first two principal components are used to create a density kernel (panel **c**). The different bandwidths presented in panel **c**, reflect the amount of data contained within each area. This kernel is then translated into the geographical space (panel **d**) and then used for the background sampling. All density kernels are calculated using a constant grid size of 1000 by 1000 and a bandwidth that corresponds to the 10% of the difference in latitudinal and longitudinal ranges of the study area.

**Model parameters and selection**

**Table S3**. Parameters returned by AutoMaxent, related to the models (mod.id) adjusted using the specified adjustments: β-values between 1-5, a random set of adjustment features, 10 thousand background points sampled using the density of presence records. **Train** and **Test AUC** values are calculated by MaxEnt, as well as the True Statistical Skill (TSS) threshold for the models (TSS MaxEnt). Since these metrics are sensitive to the threshold at which the confusion matrix is calculated, AutoMaxent also calculates its own **TSS threshold**, **mean** and **max** values, along with the continuous **Boyce** index. If AutoMaxent is set to select and filter the models, the corrected Akaike information criteria (**AICc**) of each model are calculated, along with its **Log Odds,** number of coefficients (**n-coefs**) and the difference in AICc (**Δ AICc**) with respect to the best fitted model (the model with the lowest AICc). In addition to these parameters, AutoMaxent also returns the list of MaxEnt arguments used to run the individual models, as well as the data and spatial information needed to replicate the results in MaxEnt.

| mod.id | Train AUC | Test AUC | Diff AUC | TSS MaxEnt | TSS threshold | TSS mean | TSS max | Boyce | n-coefs | Log Odds | AICc | Δ AICc |
| --- | --- | --- | --- | --- | --- | --- | --- | --- | --- | --- | --- | --- |
| 6 | 0.670 | 0.662 | 0.009 | 0.565 | 0.540 | 0.243 | 0.428 | 0.999 | 149.000 | -65914.784 | 132133.623 | 0.000 |
| 39 | 0.663 | 0.658 | 0.005 | 0.596 | 0.580 | 0.232 | 0.419 | 0.998 | 79.000 | -66302.189 | 132764.073 | 630.451 |
| 12 | 0.663 | 0.658 | 0.004 | 0.569 | 0.570 | 0.236 | 0.413 | 0.999 | 95.000 | -66297.106 | 132786.665 | 653.042 |
| 27 | 0.659 | 0.657 | 0.002 | 0.609 | 0.590 | 0.230 | 0.410 | 0.998 | 58.000 | -66511.016 | 133138.947 | 1005.325 |
| 8 | 0.658 | 0.655 | 0.002 | 0.573 | 0.560 | 0.224 | 0.398 | 0.998 | 62.000 | -66526.741 | 133178.528 | 1044.905 |
| 2 | 0.658 | 0.656 | 0.002 | 0.616 | 0.590 | 0.227 | 0.408 | 0.997 | 51.000 | -66651.152 | 133405.014 | 1271.391 |
| 40 | 0.656 | 0.654 | 0.002 | 0.576 | 0.550 | 0.220 | 0.397 | 0.994 | 43.000 | -66664.750 | 133416.005 | 1282.382 |
| 34 | 0.652 | 0.650 | 0.002 | 0.561 | 0.540 | 0.210 | 0.393 | 0.993 | 39.000 | -66692.688 | 133463.793 | 1330.171 |
| 7 | 0.652 | 0.649 | 0.002 | 0.566 | 0.550 | 0.206 | 0.392 | 0.992 | 40.000 | -66717.134 | 133514.705 | 1381.083 |
| 23 | 0.656 | 0.655 | 0.001 | 0.669 | 0.590 | 0.222 | 0.400 | 0.997 | 47.000 | -66784.333 | 133663.269 | 1529.647 |
| 3 | 0.655 | 0.654 | 0.001 | 0.598 | 0.590 | 0.222 | 0.400 | 0.999 | 48.000 | -66784.399 | 133665.427 | 1531.804 |
| 21 | 0.653 | 0.652 | 0.001 | 0.573 | 0.560 | 0.213 | 0.391 | 0.991 | 38.000 | -66799.617 | 133675.629 | 1542.006 |
| 43 | 0.650 | 0.647 | 0.003 | 0.555 | 0.520 | 0.202 | 0.390 | 0.987 | 30.000 | -66883.059 | 133826.367 | 1692.744 |
| 26 | 0.650 | 0.647 | 0.003 | 0.553 | 0.520 | 0.202 | 0.390 | 0.987 | 40.000 | -66882.605 | 133845.648 | 1712.025 |
| 17 | 0.649 | 0.646 | 0.002 | 0.557 | 0.520 | 0.203 | 0.383 | 0.992 | 32.000 | -66968.592 | 134001.465 | 1867.843 |
| 19 | 0.649 | 0.646 | 0.002 | 0.575 | 0.520 | 0.200 | 0.384 | 0.992 | 33.000 | -66967.974 | 134002.247 | 1868.624 |
| 41 | 0.630 | 0.630 | 0.000 | 0.559 | 0.560 | 0.164 | 0.359 | 0.860 | 9.000 | -67303.850 | 134625.723 | 2492.101 |
| 13 | 0.627 | 0.627 | 0.000 | 0.562 | 0.530 | 0.166 | 0.349 | 0.976 | 7.000 | -67372.246 | 134758.507 | 2624.884 |
| 22 | 0.629 | 0.628 | 0.001 | 0.571 | 0.550 | 0.151 | 0.345 | 0.753 | 8.000 | -67375.449 | 134766.916 | 2633.294 |
| 25 | 0.629 | 0.629 | 0.001 | 0.575 | 0.570 | 0.161 | 0.354 | 0.979 | 9.000 | -67399.148 | 134816.320 | 2682.697 |
| 37 | 0.627 | 0.627 | 0.000 | 0.576 | 0.530 | 0.159 | 0.349 | 0.983 | 8.000 | -67441.766 | 134899.551 | 2765.929 |
| 16 | 0.627 | 0.626 | 0.000 | 0.576 | 0.580 | 0.150 | 0.341 | 0.982 | 8.000 | -67481.053 | 134978.124 | 2844.502 |
| 11 | 0.625 | 0.625 | 0.000 | 0.563 | 0.560 | 0.162 | 0.347 | 0.978 | 3.000 | -67493.689 | 134993.382 | 2859.759 |
| 38 | 0.625 | 0.625 | 0.000 | 0.590 | 0.520 | 0.163 | 0.346 | 0.987 | 6.000 | -67537.442 | 135086.896 | 2953.273 |
| 20 | 0.625 | 0.625 | 0.000 | 0.572 | 0.570 | 0.157 | 0.349 | 0.980 | 3.000 | -67550.484 | 135106.972 | 2973.349 |
| 5 | 0.623 | 0.623 | 0.000 | 0.539 | 0.540 | 0.156 | 0.341 | 0.981 | 4.000 | -67563.413 | 135134.832 | 3001.209 |
| 9 | 0.624 | 0.624 | 0.000 | 0.577 | 0.580 | 0.153 | 0.350 | 0.980 | 3.000 | -67610.466 | 135226.934 | 3093.312 |
| 10 | 0.621 | 0.622 | -0.001 | 0.582 | 0.570 | 0.149 | 0.337 | 0.990 | 5.000 | -67663.048 | 135336.104 | 3202.481 |
| 1 | 0.624 | 0.624 | 0.000 | 0.590 | 0.570 | 0.147 | 0.352 | 0.981 | 3.000 | -67673.224 | 135352.451 | 3218.829 |

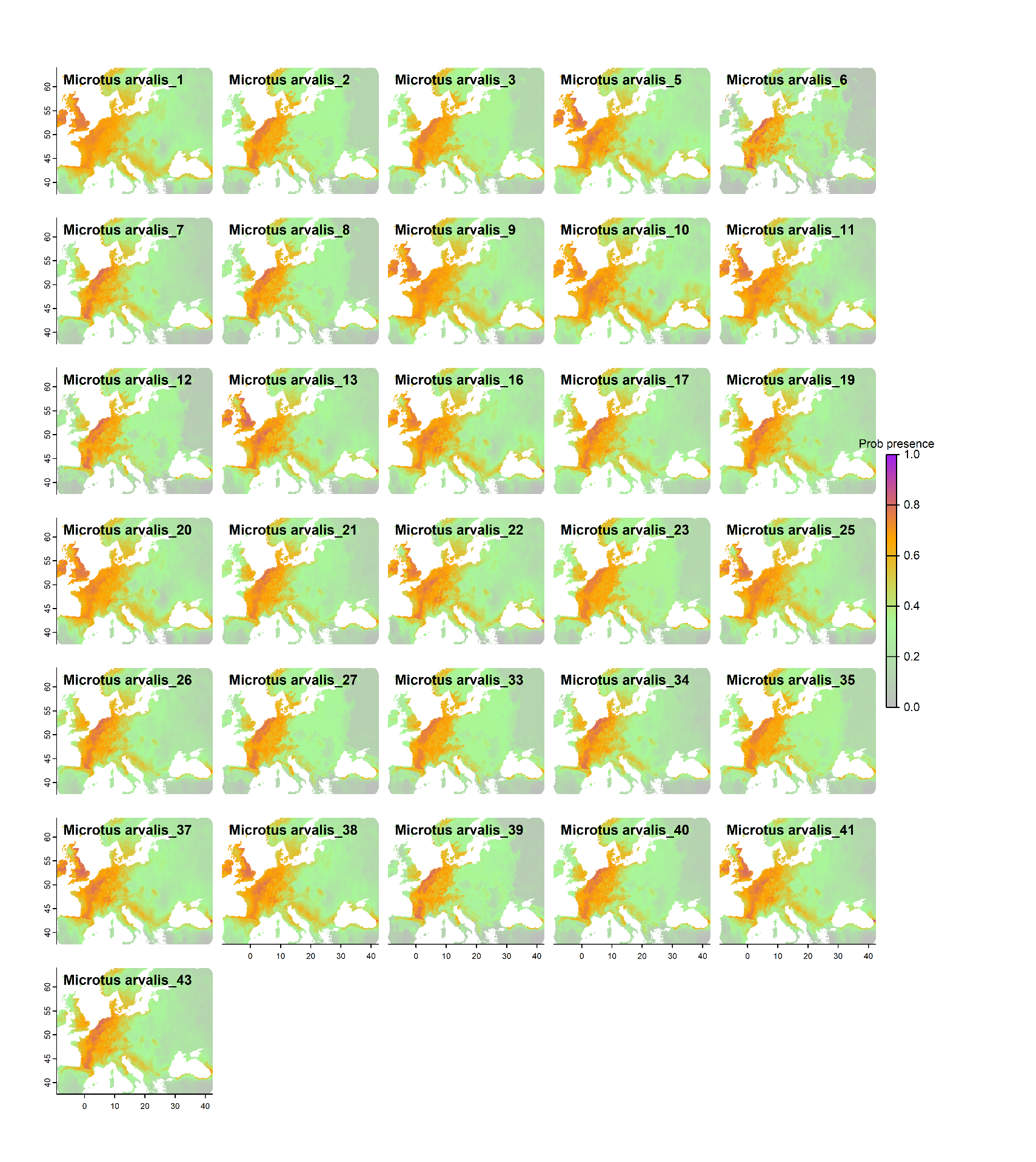

Figure S4. Predictions for the probability of occurrence of M. arvensis in Europe, for the 32 different models adjusted by AutoMaxent. Models are identified by the name of the species followed by a number, which corresponds to that of **mod.id** in **Table S3**.
